# Acute stress modulates social approach and social maintenance in adult zebrafish

**DOI:** 10.1101/2023.02.10.528021

**Authors:** Alex Cook, Holger Beckmann, Rutkay Azap, Soojin Ryu

## Abstract

Stress alters social functioning in a complex manner. An important variable determining the final effects of stress is stressor intensity. However, the precise relationship between stressor intensity and social behavior is not well understood. Here we investigate the effects of varying acute stressor intensity exposure on social behavior using adult zebrafish. We first establish a novel test using adult zebrafish that allows distinguishing fish’s drive to approach a social cue and its ability to engage and maintain social interaction within the same behavioral paradigm. Next, we combined this test with a new method to deliver an acute stress stimulus of varying intensities. Our results show that both social approach and social maintenance are reduced in adult zebrafish upon acute stress exposure in an intensity-dependent manner. Interestingly, lower stress intensity reduces social maintenance without affecting the social approach, while a higher stress level is required to alter social approach. These results provide evidence for a direct correlation between acute stressor intensity and social functioning and suggest that distinct steps in social behavior are modulated differentially by the acute stress level.

**SIGNIFICANCE STATEMENT:** Acute stress exposure has a potent effect on social behavior in many animals including humans. However, so far the effect of different stressor intensity on distinct steps of social behavior has not been directly tested. Here, using zebrafish, we develop a new social behavior paradigm and a new graded acute stressor delivery method in order to test the relationship between acute stressor intensity and social behavior. Our results show that acute stress modulates both social approach and social maintenance in an intensity-dependent manner but social maintenance is affected at lower stress intensity than social approach. Thus, this work reveals that distinct steps in social behavior are differentially modulated by the acute stress level.

## INTRODUCTION

Acute stress can profoundly impact social functioning of animals including humans (Sandi and Haller, 2015). However, the effects vary depending on the type of stressor, context, sex of the individual, specific tests employed, etc. (von Dawans et al., 2021) (Beery and Kaufer, 2015). In rodents, studies suggest that acute stress exposure leads to reduced social motivation and interaction while increasing aggression (Anacker et al., 2016; Spencer, 2017). However, other studies demonstrate an increase in affiliative behavior and reduced aggression upon acute stress exposure (Wood et al., 2003; Muroy et al., 2016). In a striking example of varied stress effects, in prairie voles, the effects of acute stress on social preferences are sexually dimorphic (DeVries et al., 1995; DeVries et al., 1996). An important variable determining the final effects of stress is the stressor intensity. Indeed, stress intensity and memory function follow the inverted U-shape in rodents (Salehi et al., 2010). Furthermore, the performance of innate behavior improves under moderate but not higher levels of acute stress in developing zebrafish larvae (Ryu and De Marco, 2017). However, the precise relationship between stressor intensity and social functioning is not well understood.

To determine the relationship between acute stressor intensity and social functioning, we utilize adult zebrafish in this study. Adult zebrafish show a high degree of sociality, where a range of social behaviors including aggression, shoaling, schooling, and mating have been well studied (Oliveira, 2013). Furthermore, the stress response systems of teleosts, including the Hypothalamo-pituitary-interrenal (HPI) axis, show a high degree of conservation with those of mammals (Wendelaar Bonga, 1997). Social behavior in adult zebrafish is typically tested either with a social preference test using a single fish approaching conspecifics or by measuring shoaling parameters within a group of fish (Ogi et al., 2021) (Miller and Gerlai, 2012). Some studies have examined the effect of acute stress on social behavior. For example, the presence of an aerial predator model or exposure to a novel environment resulted in altered group behavior, including higher shoal density in adult zebrafish (Miller and Gerlai, 2007; Kleinhappel et al., 2019). Using a social preference test, a reduction in social interaction can be induced by stress (Barcellos et al., 2020) or by pairing the social cue with an aversive stimulus (van Staden et al., 2020). However, the effect of varying stressor intensity on social behavior has not been studied so far in zebrafish.

Social functioning involves distinct behavioral sequences, broadly categorized as social recognition, social reward, and social maintenance (Chevallier et al., 2012). Several tests exist in rodents that measure different aspects of social behavior (Silverman et al., 2010). In contrast, current zebrafish social preference or shoaling tests do not allow for distinguishing different aspects of social behavior. Therefore, although social deficits in stressed fish have been reported (Abreu et al., 2016a; Giacomini et al., 2016; Barcellos et al., 2020), it is unclear which aspect of social functioning is affected. Therefore, a new behavioral paradigm is required to address this question. Typical methods to measure an animal’s motivation require animals to overcome a cost to gain access to reward (Dawkins, 1990; Kirkden and Pajor, 2006). Utilizing such a principle, here we report, a novel behavioral paradigm that measures adult zebrafish’s drive to approach social cues and their ability to engage and maintain social interaction within the same experiment. We next combine this test with a new delivery method for graded acute stress exposure using a symbolic representation of an aerial-looming predator. Mimicking the aerial predator, a computer-generated looming dot presentation is used widely in combination with activity imaging to elicit robust escape behavior and to study fear-related circuits in larval zebrafish (Temizer et al., 2015). However, although used widely in larval zebrafish and in rodents, the looming dot (LD) has not been used in adult zebrafish. Moreover, currently, there is no effective method to vary acute stressor intensity delivered to adult zebrafish. Therefore, we developed a suitable method for highly controllable and graded delivery of looming predator-related stimuli for adult fish. By combining the behavioral paradigm with graded acute stress delivery, we find that acute stress reduces both social approach and social maintenance in adult zebrafish in an intensity-dependent manner. Strikingly, we observed that social maintenance is affected by acute stressor exposure intensity that does not affect social approach. This indicates that distinct steps in social behavior may be modulated differentially by acute stress level and provides an entry point for dissecting underlying neural mechanisms.

## RESULTS

### A novel assay affords distinguishing a drive for social approach and social maintenance

To measure a drive for social interaction, we sought to create a conflict between an aversive cue and rewarding social stimuli, where fish have to overcome an aversive area in order to gain access to the reward. In our paradigm, the test fish is introduced into the main tank that has two colored zones. This tank is separated from another tank holding five conspecific fish, which only become visible when a divider positioned between these two tanks is removed (Figure 1a). Zebrafish exhibit color preference and show a preference for blue (Bault et al., 2015; Park et al., 2016). When placed in a tank with blue and white-colored areas and with the divider in place, 89.2 % of fish enter the blue zone in the first 10 seconds, after being netted into the center of the tank. The blue preference and white avoidance are maintained for a minimum of 10 minutes, during which period, the average time in the blue zone did not fall below 69% (Figure 1b). When the divider is removed after 60 seconds, revealing the social cue, fish leave the blue zone and enter the white zone, which they previously avoided. For the initial 60 seconds with the divider in place, fish spend on average 18.3% of their time in the white zone, this increases to 96.6% for the latter 120 seconds of the trial when the social cue is visible (Figure 1B). After entering the white zone, fish approach the social cue directly. The fish swims as close to the wall as possible and exhibits repetitive lateral movements along the left-right (L-R) axis followed by sharp turns to come back to the wall separating it from its conspecifics. This stereotypical movement results in a C-shaped zone within the white zone, which we term the interaction zone (Figure 1C). The average time that test fish take from first leaving the blue zone to entering the interaction zone is 2.21 seconds, indicating that fish leave the blue zone in order to approach the social cue. With the divider removed, 94.7% of fish leave the blue zone within 12 seconds to enter the white zone (Figure 1D). During the 120-second period after removing the divider, fish spend an average of 91.26 % of their time in the interaction zone in close proximity to the social cue (Figure 1E). We use females for all our behavioral tests in this study. However, the time in the interaction zone over 120 seconds were similar between female and male test fish when tested against five female social cue fish (Supplementary Figure 1). Within the interaction zone, we noted that fish make stereotypical and repetitive L-R lateral movements. We define each one of these lateral movements as one interaction regardless of the direction. One interaction can be left to right or right to left. On average a fish performs interactions at a frequency of about one interaction per second, which is consistent across the 120-second trial period (Figure 1F).

**Figure 1.**
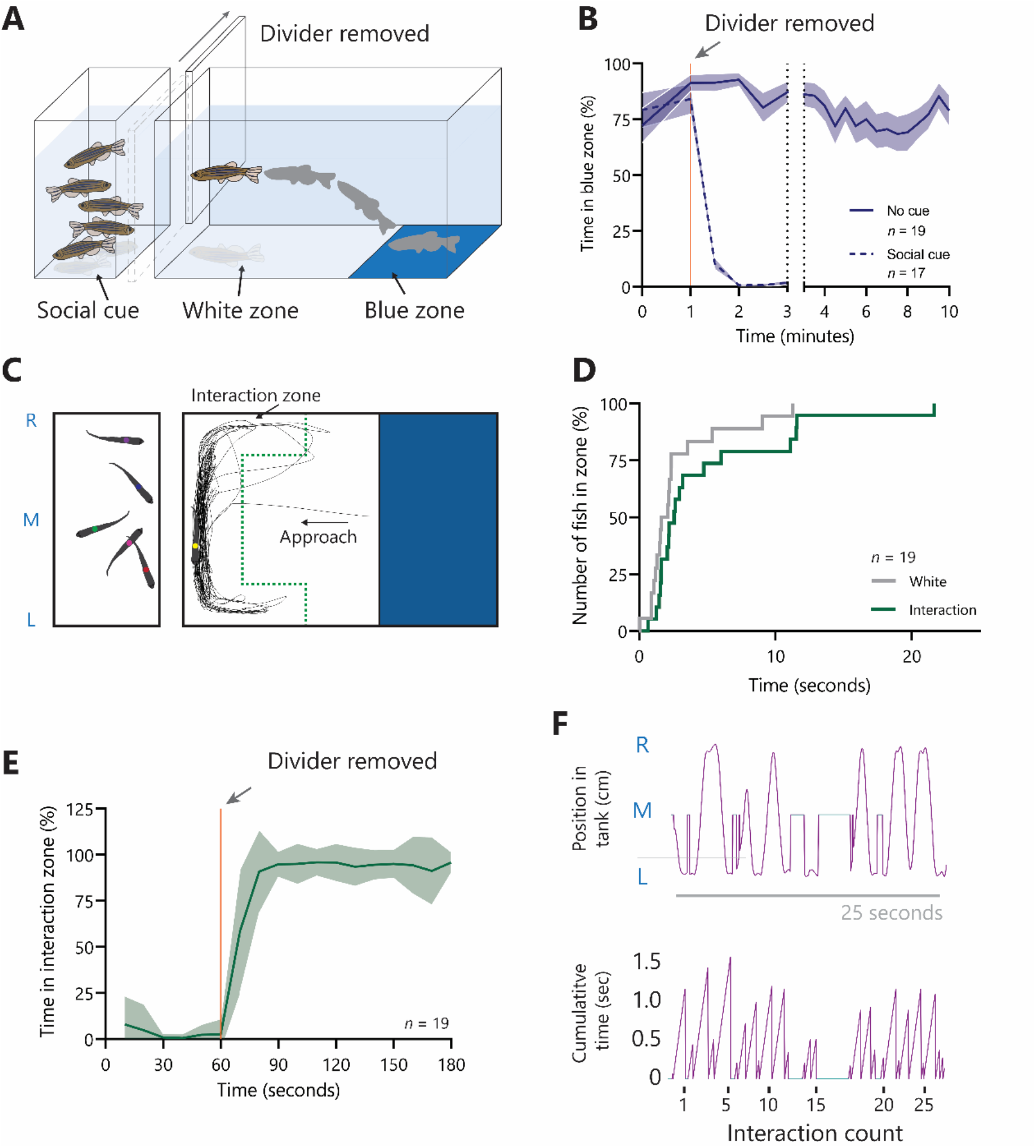
A novel social behavior assay distinguishes between social approach and social maintenance behaviors. **A.** A 3D schematic representation of the social assay experimental setup with a divider obstructing the visual social cue. **B.** Preference for the blue zone (%) with or without the divider blocking the visual access to social cues in 30 second bins. **C.** A 2D schematic representation from the top view demonstrating typical traces of the test fish. The interaction zone is marked by the green dotted line. **D.** A survival plot showing the time of the first entry of each fish to the white zone and subsequently to the interaction zone. Each event represents an individual’s first entry to each zone, displayed as a percentage of the population (%). **E.** Time spent in the interaction zone with the divider (60 seconds) or without the divider (120 seconds) as a percentage (%) of 30-second bins. F. The position of the test fish on the L-R axis during a typical 25 seconds period demonstrating stereotypical social interaction behavior. The cumulative time for each interaction is plotted demonstrating the typical duration and the frequency of interactions. Abbreviations: R, right; M, middle; L, left.

### Approach and interaction are dependent on the perceived salience of the social cue

To test whether the approach and interaction exhibited by the fish in our test represent a true social behavior rather than a novelty-related response to moving objects, we changed the salience of the cue and measured both approach and maintenance of the interaction. First, we compared different strains as the social cue, using the same strain as the test fish (TU), and two strains with different visual features. The Casper strain has a transparent appearance and the TL strain show larger fins. We then assessed the approach behavior by measuring both white zone preference and latency to approach social cues and normalized both measures to those of the TU fish (Figure 2A). We found a significant difference in the approach behavior among the three strains (Kruskall-Wallis test, H(3) = 10.18, *P* = 0.003) with a significant difference between TU and TL (** *P* = 0.004). Next, we assessed the maintenance of interaction by measuring both time in the interaction zone and the number of interactions of the three different strains (Figure 2B). A significant difference among the strains was found (one-way ANOVA, *F*(2, 18) = 23.13;*p* < 0.001) with clear differences between TU and TL (Tukey’s post hoc analysis; *** *P* < 0.001) and TL and Casper (Tukey’s post hoc analysis; *** *P* < 0.001). Next, we measured the effect of decreasing the number of conspecifics in the social cue tank. Approach was affected by the number of fish in the social cue tank (Kruskall-Wallis, H(3) = 27.03, *P* < 0.001). Fish spent more time with five fish compared to one fish (* *P* < 0.075) or when no fish (*** *P* < 0.001) were present (Figure 3C). Maintenance of interaction was also affected by the number of conspecifics (one-way ANOVA, *F*(2, 38) = 128.9; *p* < 0.001) with a reduction in maintenance with both no (*** *P* < 0.001) or one (*** *P* < 0.001) conspecific compared to five (Figure 2D). Lastly, to determine whether approach and maintenance behavior can truly represent social behavior that requires reciprocal interaction, we next assessed whether the conspecific fish used as a social cue exhibited signs of reciprocal behavior in the presence of the test fish. To this end, we measured the lateral distance (red double-headed arrow) between the test fish and the nearest social cue fish found within 1 cm of the wall separating them in order to determine whether the social cue fish also changes their relative position in order to be near to the test fish. The lateral distance between the test fish and the social cue fish on average is closer when the test fish is performing interactions compared to when it is not and in the white zone (Figure 2E). The lateral distance was compared using 258 random interaction zone distances and 243 random non-interaction zone distances (Mann Whitney test, U(501) = 25529, p < 0.001). The distances were normalized and plotted as a histogram where the test fish is 0 (Figures 2E and F). Taken together these results argue that our behavioral paradigm measures true social behavior involving both a drive to approach social cues and the ability to maintain reciprocated social interaction.

**Figure 2.**
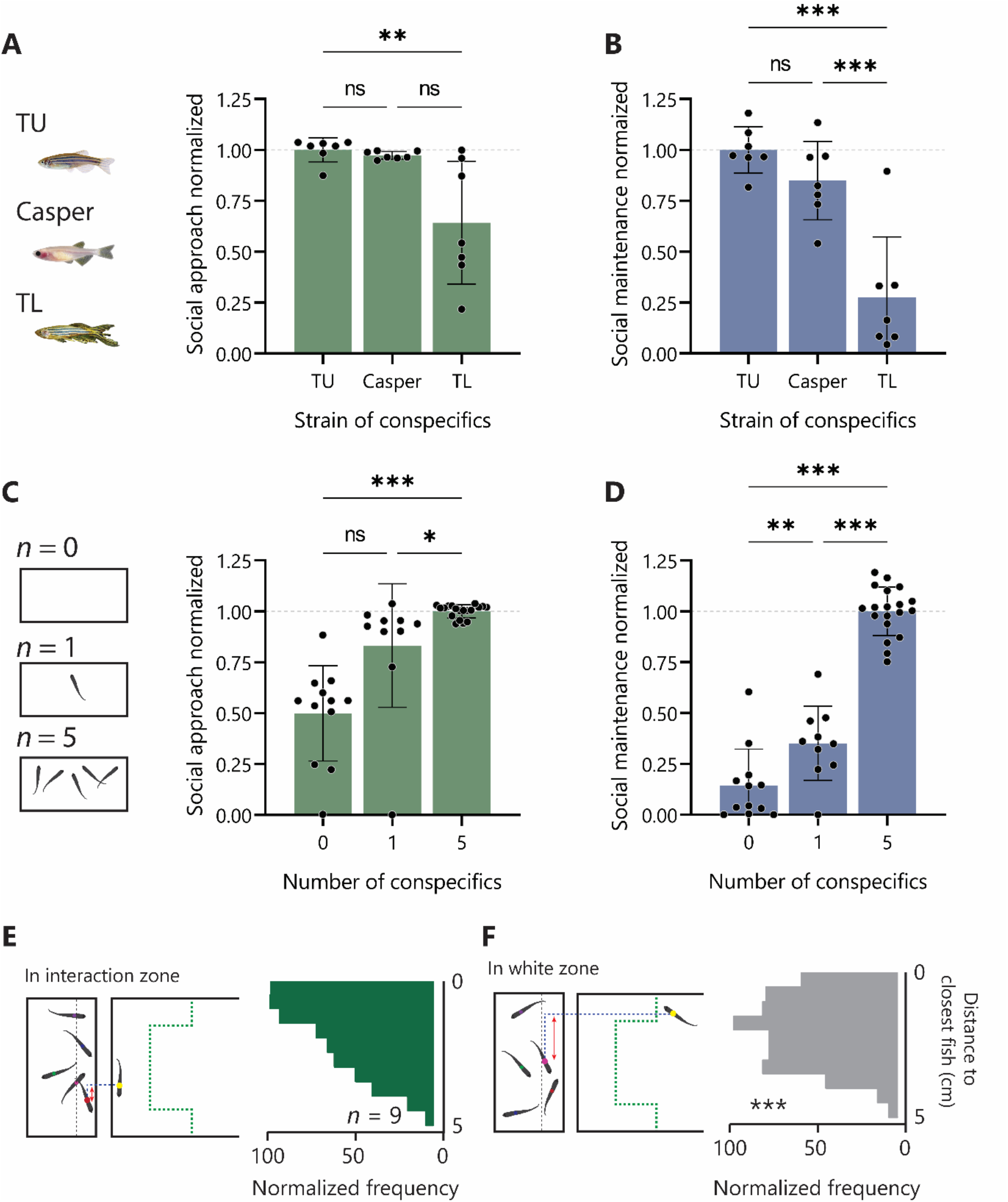
Social approach and reciprocal social maintenance behaviors are malleable to changes in the salience of the social cue. **A.** Social approach behavior (normalized white zone preference and interaction zone latency) with TU (same as test fish), Casper or TL fish as the social cue. **B.** Social maintenance behavior (normalized time in the interaction zone and interaction count) with different strains as conspecifics. **C.** Social approach behavior with 0, 1, or 5 fish as the social cue. **D.** Social maintenance behavior with different numbers of fish as the social cue. **E.** The distance to the nearest fish is measured between the test fish and the nearest fish on the L-R axis that is within the 1 cm proximity zone (grey dashed line). The distance to the nearest fish is plotted in histograms for when the test fish is in the interaction zone and when it is not in the interaction zone. Examples ‘1’ and ‘2’ are highlighted on the histogram and are illustrated above. Abbreviations: TU, tübingen; TL, Tupfel Long-fin.

### A new delivery method for inducing graded acute stress exposure in adult zebrafish

To measure the effects that acute stress has on social behavior, we designed an acute-stress paradigm that mimics an aerial-looming-predator to induce stress in adult zebrafish that consists of a small tank placed underneath a Raspberry Pi display controlled via a custom-built graphical user interface (GUI) on a master PC to generate and present graphics interchange format (GIF) images (Figure 3A). The looming dot (LD) stimulus was an expanding black circle on a white background that started at the center and filled most of the display. The custom-built GUI control panel generates a short sequence, which contains a list of 15 parameters that can be varied. We tested several parameters including the looming speed and shade of the LD but found changing the number of sequential LD that are presented, to be a highly effective means to control the intensity of the stimulus. We refer to this as LD number, which changes both the number of stimuli and the total duration of the stressor (Figure 3B). First, to measure meaningful behavioral changes associated with LD, subjects were recorded for 30 seconds to determine baseline (Pre-LD) measurements, and then whilst exposed to one of four LD intensities: 0 LD (handling control), 6 LD, 12 LD, or 18 LD (Figure 3B). During LD presentation, subjects display escape-like behavior characterized by an increase in speed. A one-way ANOVA followed by Dunnett’s multiple comparisons test revealed a significant increase in average speed during the presentation of the first 6 dots in all LD groups compared to the 0 LD group (*F*(3, 62) = 27.45; *** *p* < 0.001) (Figure 3C). Although there is no difference in average speed among 6 LD-, 12 LD- and 18 LD-exposed groups, the cumulative distance swum during the LD presentation increases relative to the number of LD as a result of an increase in the duration of the exposure. A repeated-measures one-way ANOVA of the 18 LD group following 6 LD (mean = 442.7 cm), 12 LD (mean = 712.6 cm), and 18 LD (mean = 949.7 cm) reveals a significant increase in distance moved (*F*(1.029, 10.29) = 43.19; *** *p* < 0.001) (Figure 3D). Additionally, the turn frequency increases during LD exposure from 1.09 to 1.97 turns per second during the first 6 LD compared to the pre-LD phase (two-tailed paired t-test (t(41)=8.28,*p* < 0.001) (Figure 3E). We performed linear regression and confirmed a significant correlation between the LD number and the excess distance swum (R^2^ = 0.618, F(1, 64) = 103.5, p < 0.001) (Figure 3F). During the LD exposure, fish perform an escape-like response. However, subsequently, subjects displayed intensity-dependent immobility after 6 LD, 12 LD, and 18 LD treatments. We performed linear regression and confirmed a significant correlation between the LD number and the time spent being immobile (R^2^ = 0.573, F(1, 73) = 97.94, p < 0.001). To test whether stress-induced immobility was due to physical exhaustion or anxiety-related freezing, we pretreated fish with the anxiolytic compound Diazepam for two hours before exposing them to 12 LD (Supplementary Figure 2). Diazepam pre-treatment inhibited immobility and treated fish spent less time immobile compared to the Ctrl group (t-test (t(14)=7.9, *p* < 0.001). Next, to assess whether LD exposure induced an endocrine stress response, we performed a cortisol assay to measure whole-body cortisol levels of the fish 15 minutes after being exposed to 0 LD, 6 LD, 12 LD, or 18 LD treatment. We measured a significant correlation between the LD number and whole-body cortisol (R^2^ = 0.173, F(1, 46) = 9.641, *P* = < 0.003). Collectively these results indicate that by varying the number of LD presented an increasingly higher level of acute stress exposure can be induced as evidenced by an increase in both endocrine and behavioral stress reactions.

**Fig. 3.**
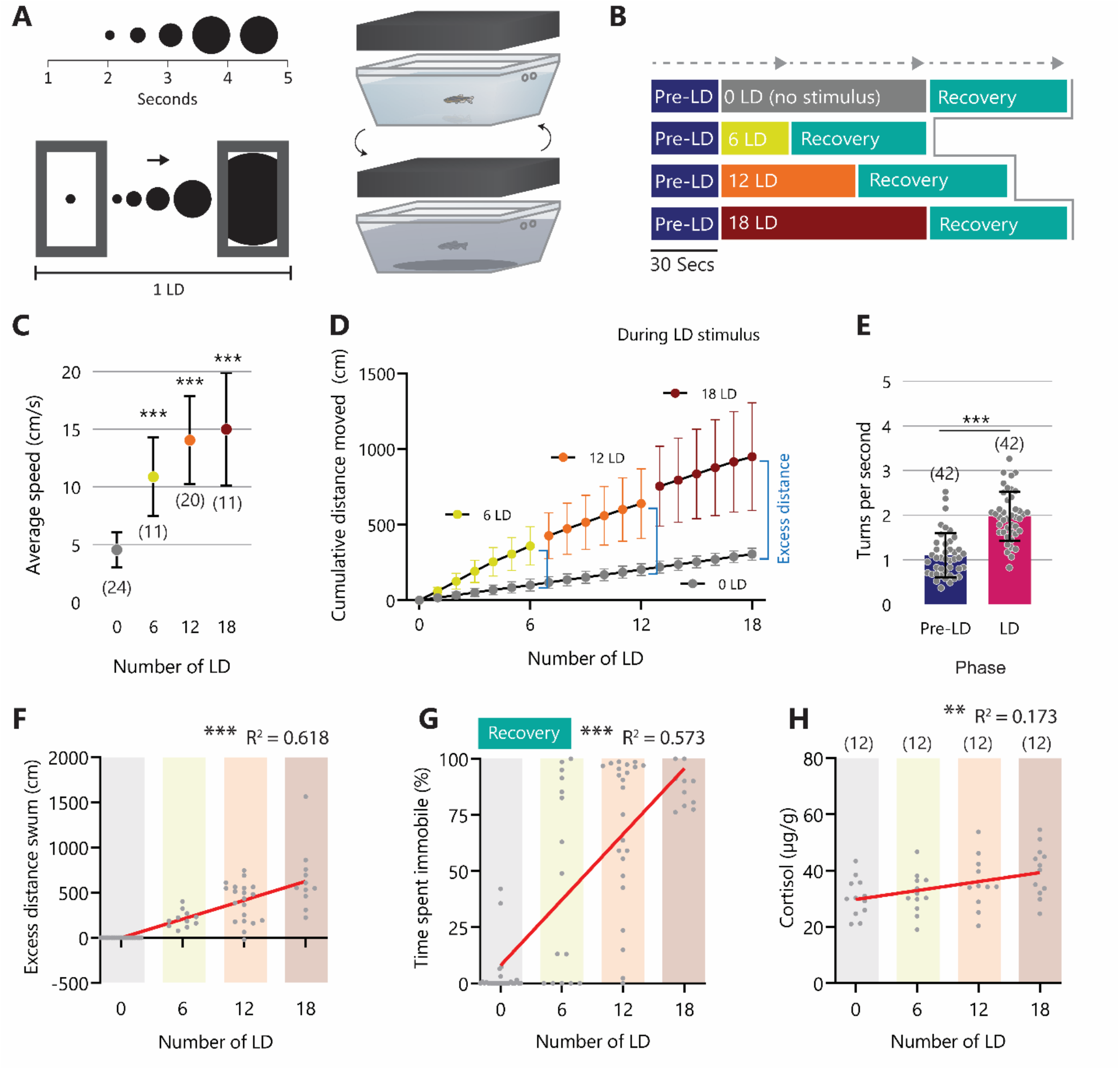
The stress intensity of LD stimulus is scalable leading to relative increases in behavioral and physiological responses. **A.** The LD stimulus is presented on a monitor above the fish and each LD event lasts 4.8 seconds. **B.** The LD is presented after a 30 seconds pre-LD phase and is then repeated 6, 12 or 18 times. A 30 seconds recovery phase is subsequently recorded. **C.** The average speed during the LD phase for each intensity (cm/s). **D.** Cumulative distance swum (cm) during the LD presentation for all LD groups. Excess distance (cm) is the delta between 0 LD and the cumulative distance at each LD number (6, 12, 18). **E.** The number of turns (turns per second) during the first 6 LD in the 6LD, 12 LD, and 18 LD group compared to the pre-LD phase. F. Correlation between excess distance swum (cm) and the number of LD presented. **G.** Correlation between time spent immobile (%) for the 60 seconds post LD exposure and the number of LD presented. **H.** Correlation between whole-body cortisol (μg/g) measured 15 minutes after LD exposure and the number of LD presented. (** *P* < 0.01; *** *P* < 0.001). Abbreviations: LD, looming dot.

### Acute stress modulates social approach and social maintenance in an intensity-dependent manner

To measure the effects of acute stress intensity on approaching social cues, we measured the time spent in the white zone and the latency to approach the social cue upon exposure to different numbers of LD exposure. The time in the white zone increases significantly when the social cue is present but the magnitude of the difference between the time spent in the white zone before and after the social cue presentation is dependent on the LD number (Figure 4A). Two-way ANOVA reveals a significant phase effect (*F*(1, 186) = 186, *P* = 0.001) and a significant interaction between phase and LD number (*F*(1, 186) = 4.06, *P* = 0.004). There is a clear correlation between the number of LD fish were exposed to and the difference in time spent in the white zone (R^2^ = 0.09, F(1, 77) = 7.696, *P* = 0.007) (Figure 4B). LD number also had a significant effect on the latency to enter the interaction zone once the social phase was initiated (Kruskall-Wallis test, H(5) = 22.26, *P* < 0.001), and there was an increase in latency in the 12 LD group (*P* < 0.001) and the 18 LD (*P* = 0,002) group compared to the basal group (Figure 4C). The ability to approach social cues requires the ability to both recognize the social cue as well as a drive to be near the social cue. To distinguish between these two possibilities, we tested whether LD exposure impacted fish’s ability to recognize its conspecific. Even after the highest intensity 18 LD exposure, the fish changed their head direction when presented with the social cue indicating that the stress exposure did not compromise their ability to recognize the social cue (Supplementary figure 3).

**Figure 4.**
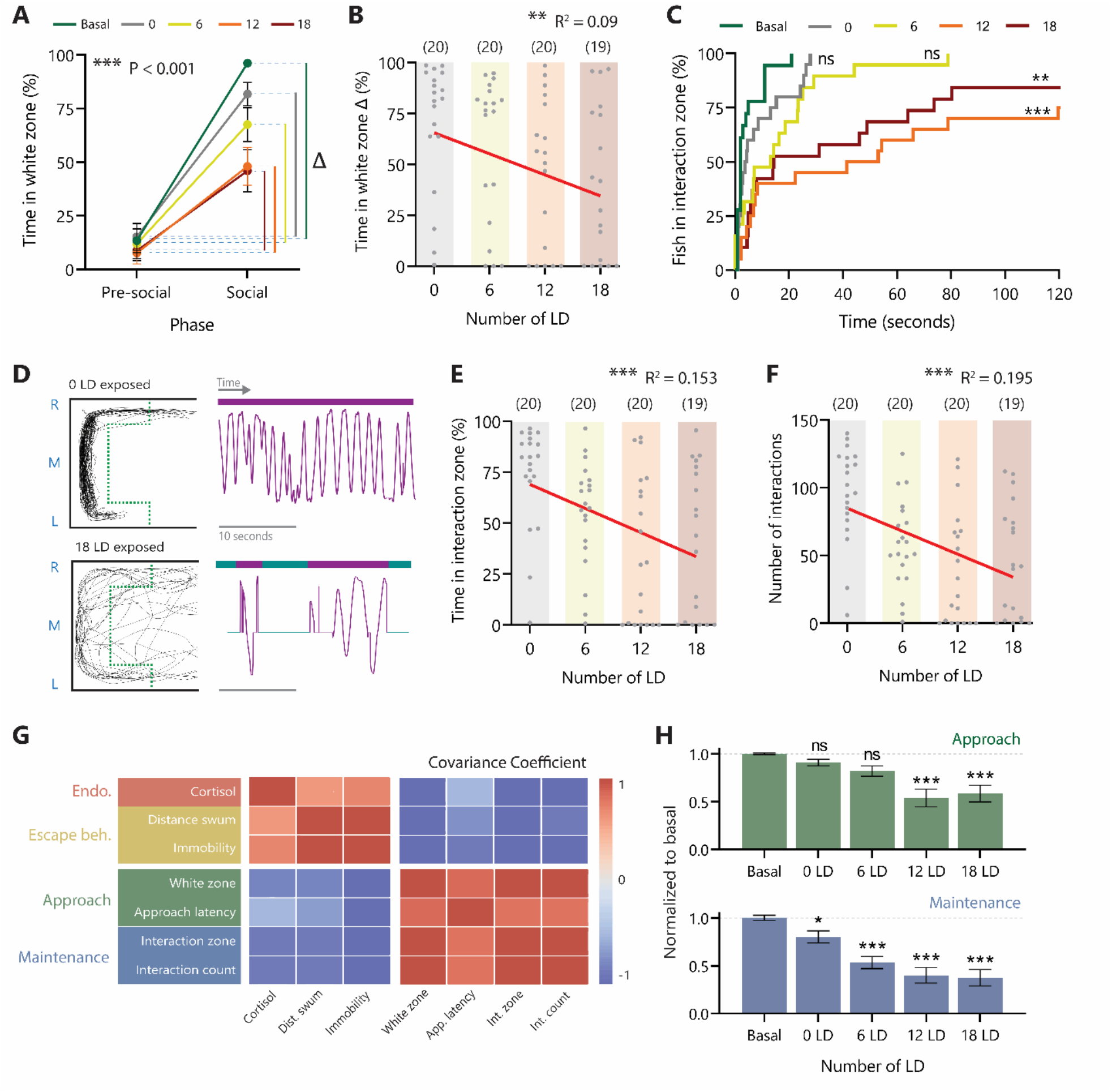
Social maintenance is modulated at a lower intensity of acute stress than social approach. **A.** Time spent in the white zone (%) with the visual divider present (pre-social) and whilst the social cue is visible (social) and the delta for each LD group. **B.** Correlation between the time in the white zone delta (%) and the number of LD presented. **C.** The number of fish (%) arrived in the interaction zone for each LD group following the removal of the divider. **D.** Example traces from above of a 0 LD and an 18 LD fish during the social phase and a 25 second example of the fish’ movement in the interaction zone on the L-R axis over time demonstrating the frequency of social interactions. **E.** Correlation between the time spent in the interaction zone (%) and the number of LD presented. **F.** Correlation between the number of interactions performed and number of LD fish were exposed to. (ns *P* > 0.05; *** *P* < 0.001). **G.** Covariance coefficient matrix for each behavior categorized into endocrinological measures, escape behaviors, social approach and social maintenance. Covariance was measured across all treatment groups. **H.** Social approach and social maintenance measures normalized to basal for each LD intensity. (ns *P* > 0.05; * *P* < 0.05; ** *P* < 0.01; *** *P* < 0.001).

Next, we measured how fish performed and maintained social interactions for the duration of the trial after being subjected to LD exposure. Example traces of a fish exposed to 0 LD or 18 LD indicate that the stereotypical ‘C’ shape pattern as well as the frequency of interactions are compromised in the 18 LD-exposed group (Figure 4D). There is a significant correlation between LD number and the time spent in the interaction zone (R^2^ = 0.153, F(1, 77) = 13.90, *P* < 0.001) (Figure 4E). Further, there is a significant correlation between the number of LD and the number of interactions performed (R^2^ = 0.195, F(1, 77) = 18.59, *P* < 0.001) (Figure 4F). Overall, as the LD number increases the endocrine and behavioral stress measures increase while the social behavior measures decrease. Endocrine and stress behavior measures have a negative covariance with both social approach and social maintenance measures across all of the treatment groups (Figure 4G). Interestingly, the extent of this negative covariance is different between the social approach and social maintenance at each treatment level. Social approach behavior is affected by LD exposure (One-way ANOVA, F(4, 92) = 9.89, *P* < 0.001), but it is only affected at higher intensities represented by 12 LD and 18 LD (Šídák’s multiple comparisons post hoc analysis for 12 LD (*P* < 0.001) and 18 LD (*P* < 0.001)). In contrast, while social maintenance is also affected by stress (One-way ANOVA, F(4, 92) = 9.89, *P* < 0.001) significant changes in behavior are observed at every treatment level (Šídák’s multiple comparisons post hoc analysis, 0 LD (*P* < 0.05), 6 LD (*P* < 0.001), 12 LD (*P* < 0.001) and 18 LD (*P* < 0.001)). This result indicates that social maintenance is affected at a lower level of acute stress exposure than social approach.

## DISCUSSION

Here we combine a novel behavior test and a new acute stress delivery method to demonstrate that acute stress modulates social approach and maintenance in an intensity-dependent manner in adult zebrafish. Strikingly our results suggest that distinct steps in social behavior are differentially susceptible to the effects of acute stress exposure.

The link between stress exposure and social behavior has been consistently documented in many species including humans. However, social behavior was measured at different time points, using different types of stressors in different contexts, which produced a complex range of outcomes. In humans, acute stress initially increases motivation to perform pro-social and social approach behavior (von Dawans et al., 2012; Buchanan and Preston, 2014). However, this effect diminishes over time (Margittai et al., 2015) and even reverses (Vinkers et al., 2013). The pro-social effect also depends on the type of stressor an individual is exposed to (von Dawans et al., 2018). Further, exposure to stress leads to a variety of outcomes in a societal context. For example, frustration that is driven by income inequality results in a more aggressive society (Wilkinson, 2004), exposure to traumatic stress is associated with social withdrawal (Davidson et al., 1991) and an increase in affiliative behavior is a common response to bullying (Kendrick et al., 2012). The complexity of outcomes is further compounded by the unique experiences, genetics, and society of the individuals observed in epidemiological studies.

Like humans, rodents exhibit social behavior that is modulated by stress and many studies have investigated the effects that different stressors have on social behavior. Often studies assessed unfamiliar male-to-male dyadic interactions, which mainly results in an increase in aggressiveness or social avoidance rather than pro-affiliative behaviors (Toth and Neumann, 2013). This outcome has been observed by both acute threat (File and Hyde, 1978; de Almeida and Miczek, 2002) and by prolonged stress administered at different developmental time points including prenatal (de Souza et al., 2013), neonatal (Wei et al., 2013), peripubertal, (Márquez et al., 2013), and adult (van der Kooij et al., 2014b; van der Kooij et al., 2014a). Activation of the hypothalamic-pituitarya-drenal (HPA) axis by corticosterone administration alone can also induce similar outcomes (Leshner and Schwartz, 1977; Mikics et al., 2004; Veenit et al., 2013). It is well established that motivation to engage in pro-social behavior is diminished in more hostile environments (File and Hyde, 1978). Therefore, when tested in a setting where the social cue is familiar and accessible, a group of freely moving familiar cage-mates exhibit pro-social behavior in the form of huddling in response to a potential threat (Kendig et al., 2011; Bowen et al., 2013).

Given the complexity of the effect of stress on social outcomes, it is critical to control relevant parameters involving both stress exposure and social behavior. A crucial variable in assessing social behavior is the use of specific behavioral assays. Social functioning involves distinct behavioral sequences and a large number of social behavior assays available in rodents emphasize different aspects of social behavior. A widely used social interaction test allows interaction of a test subject and a social cue without a physical divider or spatial restriction (File and Hyde, 1978). In contrast, the 3-chambered social approach test (Nadler et al., 2004) and the social avoidance-preference test (Berton et al., 2006) measure the choice an individual exhibits to approach a conspecific in a neutral environment and subsequent interaction with a conspecific though a physical divider in a restricted space. Social approach in these assays is usually measured as time spent in proximity to the social cue (Brodkin et al., 2004). The major difference between the way social behavior is measured in these studies compared to ours is that by including the white zone as a barrier, which needs to be overcome to initiate approaching the conspecific, we can distinguish the approach versus the interactions and the maintenance of those interactions. Social maintenance in our assay is a reflection of sustained active social behavior that goes beyond mere spatial preference, comparable to nose contacts in rodent assays (Mines et al., 2010)

Social deficits in stressed fish have been observed but differing outcomes have been reported. For example, if fish were acutely stressed and placed into an open tank with free-swimming conspecifics, an increase in shoal density is observed (Miller and Gerlai, 2007; Green et al., 2012; Kleinhappel et al., 2019). However, in a tank in which the stressed subject must approach a physically separated visual social cue, a reduction in socializing is observed (Abreu et al., 2016b; Giacomini et al., 2016; Barcellos et al., 2020). Similarly, in a three-chamber social interaction assay under aversive lighting conditions (940 LUX), zebrafish displayed no preference for the social cue, but an increase in social preference was observed following administration of the anxiolytic compound Buspirone (Barba-Escobedo and Gould, 2012). Chronic stress studies in zebrafish have led to even more divergent outcomes: during a chronic unpredictable stress protocol, subjects displayed a loosening of group cohesion after each stressor (Pavlidis et al., 2015). The same outcome was observed after 14 days of unpredictable chronic stress, but not after seven days, where the opposite was observed (Piato et al., 2011). A different type of chronic stress, early social deprivation, led to a reduction in social preference (Tunbak et al., 2020). In all of these previous investigations into the effects of stress on social behavior in zebrafish, it has been difficult to control two important parameters namely stressor intensity and specific steps in social behavior that are demonstrated and measured.

To measure a drive to approach a social cue, we created a conflict between aversive stimuli and attractive stimuli, where a subject is required to expand resources or effort to overcome an aversive cue in order to interact with the social cue. In this test, a conflict between the spontaneous avoidance of the white zone and the innate drive to approach a social cue tests the strength of the individual’s social preference. Initially, we demonstrate that the perceived salience of the social cue moderates the drive to approach and maintain proximity to the social cue. Social preference is affected by both the strain of the conspecifics and the number of conspecifics. In the less preferred scenario comprising of 1 fish or the TL strain, the innate avoidance of the white zone outweighs the drive to interact with the social cue. Next, we demonstrate how fish exposed to LD acute stress alter their social approach and maintenance behaviors. At lower intensities of stress, subjects displayed a mild reduction in approaching the social cue and entering the white zone, while at higher intensities, they largely kept away from the white zone. A consistent result was obtained in a previous study, which used the color red as an aversive cue paired with a social cue. Here, subjects initially did not approach the cue, but this trend could be reversed using dopaminergic and serotonergic manipulation (van Staden et al., 2020). A conflict could also be generated between two rewards, in which zebrafish show a less vigorous, but a more persistent tendency to approach a social reward compared to food (Daggett et al., 2019). Studies show, that similar to rodents, approaching a social cue in an unbiased environment is largely governed by motivation-associated mechanisms in zebrafish. When zebrafish approached a social cue, a significant increase in dopamine and DOPAC was measured, indicating an increase in dopaminergic activity (Saif et al., 2013). Furthermore, the ablation of dopaminergic neurons leads to social deficits similar to those observed in stressed fish (Saszik and Smith, 2018). The association between dopaminergic activity and social behavior even in unbiased environments supports that approaching a social cue is a motivation-driven, goal-directed behavior. In fact, zebrafish have been shown to be motivated by multiple stimuli (Darland and Dowling, 2001; Ninkovic et al., 2006; Braida et al., 2007). Furthermore, it has been shown that zebrafish conditioned to self-administer opioids continue to self-administer throughout a progressive ratio schedule, demonstrating that not only are zebrafish able to prefer one stimulus over another but actively work in order to receive rewards (Bossé and Peterson, 2017).

In conclusion, our results demonstrate that acute stress reduces both social approach and social maintenance in adult zebrafish. Strikingly, lower stress intensity reduces social maintenance without affecting the social approach, while a higher stress level is required to alter social approach. These findings provide evidence that acute stress modulates distinct steps in social functioning differentially in an intensity-dependent manner in adult zebrafish. Determining the effects of different levels of acute stress exposure on distinct steps of social behavior is an important prerequisite in identifying correct neural correlates of how stress modifies social behavior.

## METHODS

### Animal housing, husbandry, and experiments

Tübingen (TU) wild-type adult zebrafish (*Danio rerio*) bred in our facility were raised in mixed sex groups of 40 in 11L tanks with a 12 h light (300-800 LUX - water surface)/12 h dark cycle. Fish were fed twice per day, once with artemia (Sanders, Great Salt Lake Artemia) and once with fish flakes (TetraMin Flakes). Water conditions were kept at 28 ± 0.5 °C, 550 ± 50 μS conductivity, and 7.8 ± 0.3 pH. Between six- and nine-month-old females (unless stated otherwise) (300-600 mg body weight) were separated from their home tanks and were moved into an experiment room one week before experimentation. In the experiment room, subjects were housed in groups of ten in 3 L tanks under the same housing conditions as the main housing facility. On the day of testing, fish were used in only one behavior test and all behavior experiments were carried out between 09:00 and 13:00 before feeding. 15 minutes after testing, some fish were sacrificed in ice water for cortisol extraction. Zebrafish experimental procedures were carried out in compliance with the ethical guidelines of the German animal welfare law and approved by the local government (Landesuntersuchugsamt Rheinland-Pfalz, Germany - 23 177-07/G20-1-033).

### Social assay

The social behavioral assay setup is composed of a transparent cuboidal acrylic tank measuring 18 cm (L) x 11 cm (W) x 9 cm (D) filled up to 6 cm with fish housing water (1.2 liters). The floor of the tank is covered by white vinyl (66%) and blue vinyl (33%). At the end of the white area of the tank, a white opaque acrylic divider is positioned in the 2 cm gap between the blue/white tank and the social cue tank and functions as a visual barrier. The social cue tank is a custom-made transparent cuboidal acrylic tank measuring 6 cm (L) x 13.5 cm (W) x 9 cm (D) filled up to 6 cm with fish housing water (0.7 liters) and contains five conspecific wild-type female fish who were habituated in the tank for 30 minutes before testing. The two tanks were enclosed by white panels on two sides and an infrared back panel positioned behind the test area, which was used to generate contrast between the subject and the background. A photography dome covers the arena from external light and movement. A halo light provided homogeneous ambient light at 550 LUX (at the water surface level) and 3200 Kelvin. The assay was recorded using two cameras, one positioned above and one to the side of the tank (using an infrared filter) providing 3D X, Y, Z location coordinates. An additional camera was positioned above the social cue tank to track their positions. Subjects were netted into the center of the tank and after five seconds, the recording started. After 60 seconds, the visual divider was manually moved to reveal the social cue. The subject was recorded for a further 120 seconds.

For analysis, the tank was divided into three zones: blue zone, white zone, and interaction zone (Supplementary Fig.3A, C). The interaction zone is the shape of a ‘C’; the size and shape of the zone were calculated to encompass 95% of activity of a wild-type control fish. The time spent in each zone was calculated as the percentage of the pre-social phase (first 60 seconds, no social cue visible) and the social phase (latter 120 seconds, with social cue visible) that was spent in each zone. Latency to enter a zone was measured as the initial entry to a zone following the beginning of the assay or removal of the divider (dependent on the phase stated). Interactions were counted as lateral movements in one direction whilst in the interaction zone, a new interaction started if the subject changed direction or left and reentered the interaction zone.

Social behavior is divided into two constructs, social approach and social maintenance. Social approach is the average normalized score for the time spent in the white zone (%) and the latency to approach the interaction zone (s). Social maintenance is an average normalized score for the time spent in the interaction zone (%) and the number of interactions performed.

### Looming dot acute stress stimulus

A single fish was netted from their home tank in the experiment room to a smaller tank 16.9 — 18.7 cm (L) x 8.7 cm (W) x 10 cm (H) of which three sides and the floor are sanded to reduce reflections. The tank was immediately placed underneath a Raspberry Pi display (Raspberry Pi 3 7” display 15.4 cm x 8.6 cm), mounted 2 cm from the top of the tank. Each subject was given 60 seconds to habituate to the tank before the recording began. Recording (Basler GenICam 2) and live-tracking (EthoVision XT 13, Noldus Information Technology) were performed at 30 FPS and a resolution of 1280 x 960 during the trial. After 30 seconds of recording baseline behavior (Pre-LD phase), the LD stimulus was presented on the display.

The display was controlled by a Raspberry Pi (Model B) running a Linux operating system controlled via a custom-built graphical user interface (GUI) on the master PC to generate and present graphics interchange format (GIF) images. The custom-built GUI control panel generates a short sequence, which contains a list of 15 parameters for the programs on the Raspberry PIs. This sequence can be printed as a txt file on the Raspberry PIs via WLAN (PLINK). Custom-built scripts read the txt file, which was printed by the control panel on the Raspberry PI and the visualization process starts and creates the background in the right color and the expanding dots. The looming dot (LD) stimulus was an expanding black circle on a white background that started at the center point and filled most of the display. The number of dots shown was 0, 6, 12, or 18 depending on the treatment group. The color of the dot was black on a white background, and each dot expanded for 2.8 seconds. After reaching the maximum size, the dot covered the screen for 1 second, after which a white background was presented for 1 second before the next dot was presented. Each LD event took 4.8 seconds, which means that the duration of the LD phase was equal to the number of LD x 4.8 seconds. Following the final LD presentation, the fish were recorded for a further 60 seconds (Recovery phase).

X and Y coordinates were generated from the live tracking and the data were processed in Python to generate behavior parameter data which was graphed and statistically analyzed in Prism 9 (GraphPad Software, Inc.). The average speed was calculated as the distance moved per second based on X and Y coordinates for the duration of each phase. Excess distance was the cumulative distance swum during the LD phase minus the distance swum by the 0 LD group. Turn frequency was calculated as the number of times the swimming direction changed on the X-axis per second for the duration of each phase. When combined with the social assay, fish were first subjected to the LD protocol with the desired number of LD and immediately following the recovery phase they were netted from the LD treatment tank to the social assay tank.

### Analysis of behavior

Behavior was tracked using EthoVision XT 13, Noldus Information Technology live tracking. Centroid X, Y and Z coordinates (only X and Y for LD assay) scaled to cm for each frame (30 FPS) were imported into python and converted to ‘pandas’ data frames. For the social assay Y coordinates from the top camera were converted to Z coordinates and were merged with the X and Y coordinates of the side view camera. Data from each experimental trial were concatenated and labeled by experiment date, treatment group, and fish ID number. Based on X, Y, and Z coordinates and the frame number, we calculated distance, direction and spatial metrics. Speed was calculated as distance moved (cm) per frame (cm per f) and adjusted to distance per second (cm per second = (cm per f * 30)). Freezing duration was calculated as the number of frames under the freezing threshold. The freezing threshold was set at 10 consecutive frames where less than 0.1 cm was moved. Turns were calculated as changes in direction frame-by-frame using only X and Z coordinates in the social assay and X and Y coordinates in the LD assay. Information on zone preference and entry was calculated by assigning each frame to a zone based on coordinates.

For analysis of the conspecifics, videos were generated using a top view camera. Videos were imported into python and using a script inspired by and using tools from Traktor (Sridhar et al., 2019), Centroid X and Y coordinates were generated for each conspecific, however the individual ID of each fish was not maintained. The nearest fish distance (distance on Z axis between test fish and nearest conspecific fish) was calculated based on the coordinates of the conspecific fish closest to the test fish on the Z axis that was within 1 cm of the dividing glass.

The covariance matrix was generated in python using the StandardScaler and decomposition.PCA tools by scikit-learn and plotted using the heatmap tool by seaborn. For social approach and maintenance, features were min-max normalized between 0 and the average basal score.

### Diazepam treatment

Subjects treated with diazepam were housed in groups of 10 in their home tank, which was disconnected from the circulating water system in the experiment room. Diazepam (Diazepam-ratiopharm, PZN: 02232507; Diazepam (10 mg/ml), ethanol (105 mg/ml), benzyl alcohol (30 mg/2 ml), benzoic acid (1 mg/ml), sodium benzoate (49 mg/ml), propylene glycol (400 mg/ml)) mixed in 10ml of distilled water was added to the tank to reach the desired concentration (156 nM) in a volume of 1.5L. After two hours of Diazepam treatment, subjects were placed into the LD stimulus tank for testing.

### Whole-body cortisol extraction and assay

After LD exposure, fish were allowed to rest for 15 minutes before being netted into ice water (2 °C) for 30 seconds and then they were immediately placed into a 5 ml tube and submerged into dry ice and ethanol for 10 minutes and finally frozen at −80°C until further processing. For cortisol extraction, 1 ml/100 mg (fish weight) of distilled water was added to the tube and the fish were thawed at room temperature. The fish were homogenized (25K, IKA disperser) for 60 seconds or until it was possible to pipette the homogenate. 1 ml of homogenate was moved into a 1.5 ml Eppendorf tube and centrifuged (10K RPM, 5 minutes, 4°C). The supernatant was added to an Eppendorf tube containing ethyl acetate 99.5% and vortexed for 60 seconds. The tube was centrifuged (5K RPM, 5 minutes, 4°C) again and then placed into a −80°C freezer for 10 minutes. The ethyl acetate (supernatant) was transferred to a new tube and was evaporated (Eppendorf, Concentrator 5301, 30°C). The pellet was then resuspended in 20 μl of diluent. The samples then entered a competitive cortisol assay (Cisbio HTRF^®^ Cortisol Kit, 62CRTPEG) and the plate was read (TECAN, infinite M1000 PRO). The data is calculated and presented as cortisol (μg) per weight of fish (g).

### Experimental Design and Statistical Analysis

Group differences were analyzed using GraphPad Prism 9. The *n* for each group is reported in the figure or figure legend. The variance in data for each group is represented by the SEM and individual data points are plotted when necessary. No outliers were purposefully removed during conducting the experiment or in analysis. Variations in *n* are a result of data lost during acquisition or experimental design. Control groups and certain treatment groups may have more *n* in certain experiments as baseline behaviors and effects were assessed before entire data sets were generated in order to reduce the number of animals used. In cases where a figure includes multiple experiment days, each group was tested on every experimental day and day-to-day variations were assessed and no significant differences were found.

The statistical test used for each comparison is reported in the results section. The tests were selected based on the number of comparison and distribution of the data. For comparison of two groups either an unpaired t-test (if data is gaussian) or a Mann-Whitney (non-gaussian data) test was used based of the distribution of data. In the case of the groups being the same individuals from different time points, a paired t-test was selected. For comparisons of more than two groups (data is gaussian) under the effect of one independent variable, a one-way ANOVA was used followed by either a Tukey’s post hoc analysis for comparing all possible group combinations or a Dunnett’s test for comparing all groups against one control group. For the comparison of more than two groups (data is not gaussian) and the effect of one independent variable, a Kruskall-Wallis test was employed followed by a Dunn’s post hoc test to compare groups. For assessing the effects of two independent variables (e.g., phase X treatment), a two-way ANOVA (data is gaussian) test was selected. For multiple comparisons analysis following a two-way ANOVA, a Šídák’s multiple comparisons post hoc analysis was used. Correlation is presented as the R^2^ value generated using simple linear regression analysis using GraphPad Prism. P-value adjustments were set as the default method for each multiple comparisons test in GraphPad Prism. P-values presented at 95% confidence levels with an alpha level of 0.05; ns P >= 0.05; * P < 0.05; ** P < 0.01; *** P < 0.001.

## AUTHOR CONTRIBUTIONS

HB, AC and SR Designed Research; HB and AC Performed Research; RA Contributed unpublished analytic tools, HB and AC Analyzed Data; AC and SR Wrote the paper.

## ACKNOWLEDGEMENTS

The authors would like to thank Kathrin Domdera for expert zebrafish maintenance, Christian Moh for the support in establishing Raspberry Pi setup, Fabian Rose for carrying out initial testing of LD, and Dr Rodrigo De Marco for helpful discussion and comments on the manuscript. This work was supported by the Boehringer Ingelheim Foundation; German federal Office for Education and Research (BMBF) – 01GQ1404; Dennis and Mireille Gillings Foundation.

---

**Supplementary figure 1.**
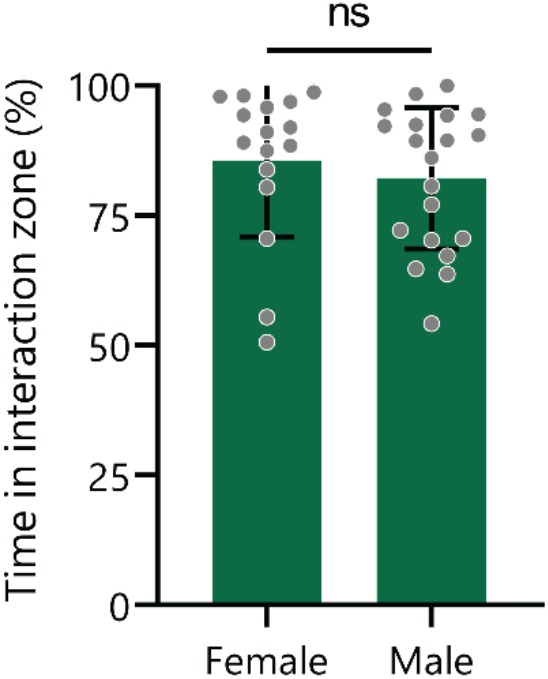
Male and female adult zebrafish display social interaction. Time in the interaction zone (%) over 120 seconds for both female and male test fish with 5 female social cue fish. (ns *P* > 0.05).

**Supplementary figure 2.**
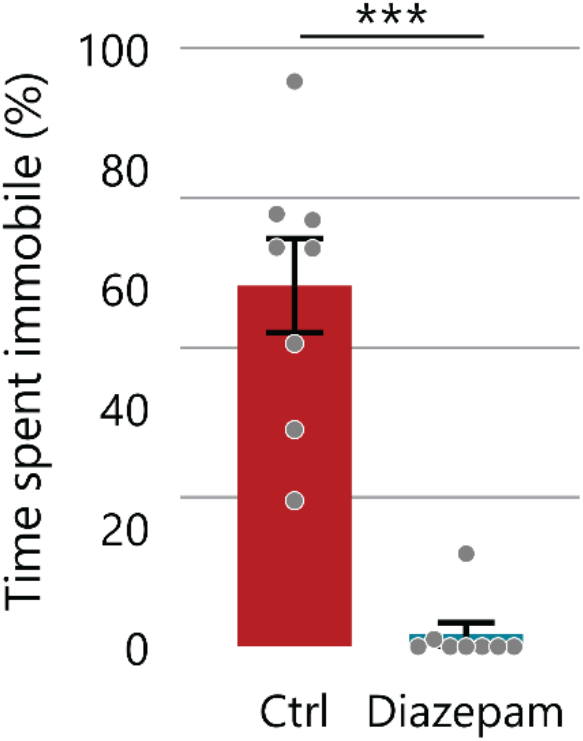
Stress induced immobility is inhibited by pre-treatment with diazepam. Time spent immobile (%) after exposure to the looming dot acute stress stimulus (12 LD) with (Diazepam) and without (Ctrl) pre-exposure to Diazepam (156 nM) for 2 hours. (*** *P* < 0.001).

**Supplementary figure 3.**
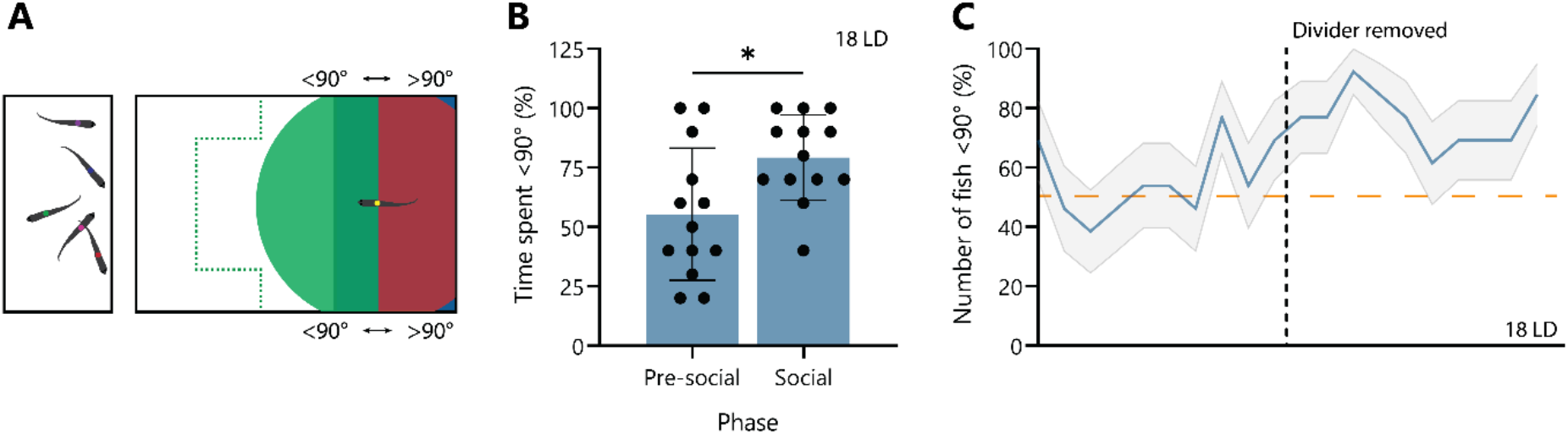
After being exposure to high intensity stress, fish still orientate towards the social cue. **A.** Heading direction of the test fish measured as >90° or <90°. **B.** Time spent <90° (facing social cue) for 18 LD exposed fish in the pre-social phase and social phase. **C.** Number of fish in the <90° direction following 18 LD exposure in the last 10 seconds of the presocial phase and the social phase. (* *P* < 0.05).

**Supplementary figure 4.**
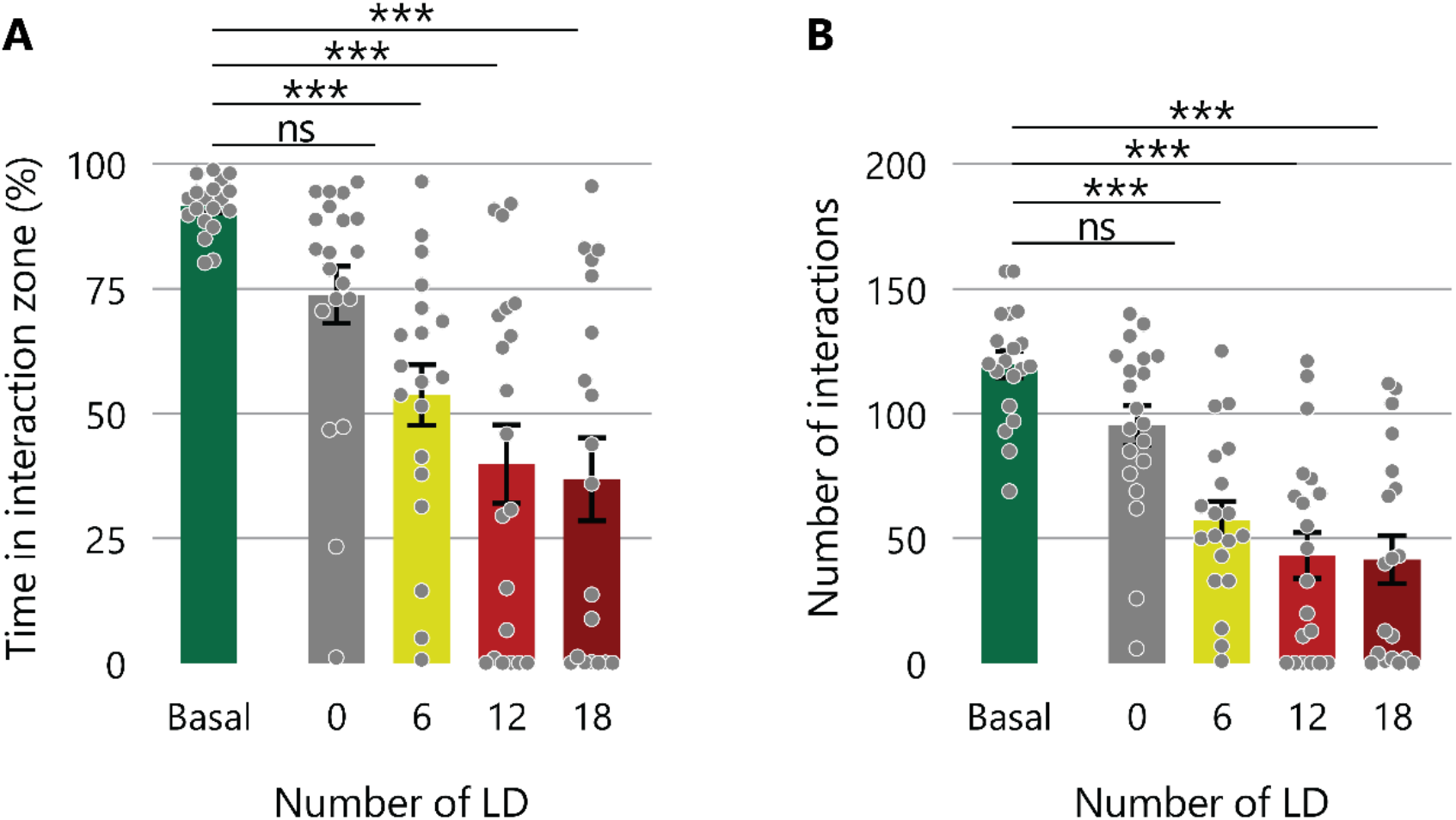
All LD-exposed groups exhibit reductions in social approach and maintenance compared to untreated fish. **A.** Time in the interaction zone (%) over 120 seconds for each treatment group. **B.** The number of interaction performed in the interaction zone during the 120 second social phase. (ns *P* > 0.05; *** *P* < 0.001). LD exposure had a clear effect on the time spent in the interaction zone (Kruskall-Wallis, H(5) = 42.38, P < 0.001) with 6 LD, 12 LD and 18 LD showing reduction compared to the basal group (Dunn’s multiple comparisons, (P < 0,001) for each of the three comparisons (Supplementary Figure 3A). The number of interactions performed also was affected by LD exposure (One-way ANOVA, F(4, 92) = 8.2, P < 0.001) with 6 LD, 12 LD and 18 LD groups all performing fewer interactions compared to the Basal group (Dunnett’s multiple comparisons test, (P < 0,001) for each of the three comparisons) (Supplementary Figure 3B).

